# The nuclear RNAi factor, NRDE2, prevents the accumulation of DNA damage during mitosis in stressful growth conditions

**DOI:** 10.1101/428250

**Authors:** Aarati Asundi, Srivats Venkataramanan, Gina Caldas Cuellar, Atsushi Suzuki, Stephen N. Floor, Andrei Goga, Noelle L’Etoile

## Abstract

Organisms have evolved multiple mechanisms to prevent and repair DNA damage to protect the integrity of the genome, particularly under stressful conditions. Unrepaired DNA damage leads to genomic instability, aneuploidy, and an increased risk for cancer. Before the cell can divide, it must repair damaged DNA and it is thought that this process requires global silencing of most transcription. In *C. elegans*, NRDE-2, in complex with other nuclear factors and guided by small RNA, directs heterochromatin formation and transcriptional silencing of targeted genes. Additionally, when *C. elegans* are cultivated at high temperatures, NRDE-2 is required to maintain germ line immortality. However, the role of NRDE-2 in maintaining the physical integrity of the genome is not understood. We show here that loss of NRDE2 in either nematode or human cells induces the accumulation of DNA damage specifically under conditions of stress, such as cultivation at a high temperature in *C. elegans* or Aurora B Kinase oncogenic overexpression in the MCF10A epithelial breast cell line. In addition, we found that NRDE2 interacts with β-actin in unstressed mammalian cells. This interaction is dramatically reduced upon DNA damage. Monomeric nuclear actin binds to heterochromatin remodeling factors and transcriptional activators while filamentous actin has been implicated in DNA repair processes. Thus, NRDE2 may dissociate from actin when it becomes filamentous as a result of DNA damage. In this way, heterochromatin factors may associate with the actin dependent DNA repair process to allow appropriate mitotic progression and maintain genomic integrity.

## Introduction

Organisms have developed multiple ways to protect genomic integrity during mitotic cell division. Maintenance of the physical and informational integrity of the genome ensures that cells accurately transmit genetic information from one generation to the next, thereby safeguarding against diseases associated with genomic instability, such as cancer. Genomic instability is caused primarily through DNA damage. DNA damage can be incurred in a number of ways, including naturally during DNA replication, meiotic recombination, improper chromosome segregation, and transposable elements. DNA damage can also be caused by external sources such as UV irradiation or drugs. If the cell attempts to divide before the DNA is repaired, the unrepaired DNA damage can result in mitotic delay, aneuploidy, or apoptosis, which in turn can lead to disorders in the organism including cancer and infertility.

To protect against genomic instability, cells must fulfill a checkpoint at the Gap2 to Mitosis (G2/M) transition [46] at which the cell repairs any DNA damage sustained during the previous cell cycle phases. If the requirements of the G2/M checkpoint are not satisfied, the cell will express a number of cell cycle inhibitors, leading to an arrest in G2. [30]

DNA repair is especially important in the germ line where strong evolutionary pressure to maintain the integrity of the genome of germ cells selects against harmful mutations. Thus, the germ line provides a window into how the environment affects the cell cycle. The nematode *Caenorhabditis elegans* has a well-characterized germ line and reproductive cycle, which can be used to study the effects of improper mitosis on cell fate and fertility. The adult *C. elegans* gonad contains a stem cell niche, which mitotically proliferates to self-renew. These proliferating germ cells (PGCs) constitute the only pool of mitotically dividing cells in the adult worm. As the PGCs divide, they travel proximally along the gonad, where they transition from mitotic proliferation to meiotic division and ultimately differentiate into spermatozoa or oocytes. Therefore, maintaining mitotic integrity in the PGC pool is essential for ensuring the reproductive integrity of the animal.

Previous studies show that the number of PGCs is susceptible to internal molecular changes as well as external stresses, which can impact the worm’s fertility. In addition to canonical signaling, such as the GLP-1/Notch, [31–32] DAF-7/TGFb [33] and IGF-1/insulin [34] pathways, many small RNA pathway proteins have also been extensively studied in the context of maintaining the PGC pool and fertility. For example, mutations in CSR-1 (argonaute that binds endogenous 22G siRNAs)[36], EGO-1 (putative RdRP required for small RNA biosynthesis) [37, 38], DCR-1 (cleaves dsRNA) [39, 40] and EKL-1 (Tudor domain protein required in 22G RNA biosynthesis) [36, 41] have all been shown to have significant mitotic and meiotic germ cell defects. Furthermore, external stresses such as starvation or changes in cultivation temperature can directly impact the PGC pool and subsequently affect fertility. [34, 35]

In particular, the nuclear RNAi protein, NRDE-2 is required to maintain reproductive integrity of *C. elegans* in response to environmental stress. *Nrde-2* null mutants remain fertile indefinitely at when grown under normal conditions at 20°C, however, when propagated at 25°C they produce fewer progeny each successive generation until becoming completely sterile by the fourth generation. [1, 16] This progressive sterility is termed a mortal germ line phenotype (Mrt). *Nrde-2* null mutant worms grown at 25°C also express high levels of repetitive RNA as compared to wild type worms propagated under the same conditions. This is consistent with the hypothesis that NRDE-2 acts to silence transposons and repress repetitive sequences, [16, 47] which are naturally occurring DNA damaging agents and can affect the informational and physical integrity of the genome.

We and others have hypothesized that the Mrt phenotype seen in *C. elegans* mutants lacking heterochromatin factors, piRNA, or nuclear RNAi factors, including NRDE-2, might result from accumulated DNA damage in the germ line. [47] The function of NRDE-2 in preventing DNA damage accumulation seems to be conserved across species but had not been directly studied in the *C. elegans* germ line. In the fission yeast, *S. pombe*, loss of the NRDE-2 homolog (known as nrl1) results in increased DNA damage, which was posited to arise from unresolved R-loops. [17] R-loops are RNA-DNA hybrids that arise from transcription of repetitive elements in which the RNA message invades the double stranded DNA helix. The DNA-RNA hybrid is more stable than the DNA-DNA hybrid and can cause DNA damage when the replication fork collides with it.

Transcriptional silencing of repetitive elements by heterochromatin factors such as NRDE2 was thought to prevent the possibility of R-loop formation. [48] A recent study in HEK293 (human embryonic kidney) cells, however, showed the human homolog of NRDE-2 (known as NRDE2 or C14orf102) plays a role in DNA damage response and preventing the accumulation of double stranded breaks, independently of R-loop formation and resolution. [22] Thus, NRDE2 may act in its capacity as a heterochromatin factor to repress transcription of repetitive elements to limit both R-loop formation and transposition, which are two events that pose threats to the integrity of the genome. Recently, it has been increasingly appreciated that heterochromatic regions of the genome (nuclear speckles) are associated with monomeric actin and that heterochromatin remodelers such as SWI/SNF, SWR1 and INO80 are tightly bound to actin monomers and this association is important for their ability to bind to chromatin. [45] Thus, stressors such as high temperature and DNA damage have been shown to tilt the balance of nuclear actin from monomeric to filamentous, which has been shown to be required for efficient DNA damage repair. [25] How heterochromatin associated factors and nuclear actin polymerization status are linked, however, remains unclear. In this paper, we show evidence that NRDE2 prevents the accumulation of DNA damage particularly under conditions of stress in proliferating cell populations. In the *C. elegans* germ line, NRDE-2 loss resulted in the accumulation of DNA damage only when the worms were propagated at a high temperature. This correlated with a decrease in the number of PGCs and was consistent with the previously reported Mrt phenotype. Moreover, in human cells, we found that NRDE2 loss did not significantly impact the proliferation of normal epithelial breast cells MCF10A unless they overexpress the oncogene Aurora B Kinase (AuBK). In these cells, we observed a delay in mitotic prophase as well as an increase in DNA damage. This indicates that NRDE2 may have a role protecting against DNA damage accumulation. Finally, we observed a strong protein-protein interaction between NRDE2 and β-actin in normally proliferating MCF10A cells, which was weakened upon the introduction of DNA damaging agents. Thus, we suggest a novel link between this heterochromatin factor, actin, and DNA repair.

## Results

### Loss of NRDE-2 coupled with propagation at high temperature incurs DNA damage in the mitotic zone of the C. elegans gonad

Previous studies in *C. elegans* have shown that loss of NRDE-2 does not affect germ line immortality when the worms are propagated at 20°C. We have observed that although *nrde-2 (gg91),* null mutants remain fertile indefinitely when cultivated at 20°C, these worms have a lower brood size as compared to wild type (Fig S1, p=0.0148, n=4). In contrast, if the worms are cultivated at 25°C, a more stressful temperature, *nrde-2 (gg91)* mutants exhibit a Mrt phenotype that increases in severity with each successive generation.[1] We asked whether the decrease in brood size of *nrde-2 (gg91)* mutant worms was a result of physiological defects in the gonad. We observed a significant decrease in the number of proliferating germ cells (PGCs) in the mitotic zone in *nrde-2 (gg91)* worms propagated at 20°C as compared to wild type (Fig 1A and B, left, p=0.0115). When we propagated the worms at the stressful temperature of 25°C for 2 generations (G2), we found that the number of PGCs was significantly decreased as compared to wild type G2 worms grown at 25°C (Fig 1A and B, left, p<0.0001).

**Figure 1.**
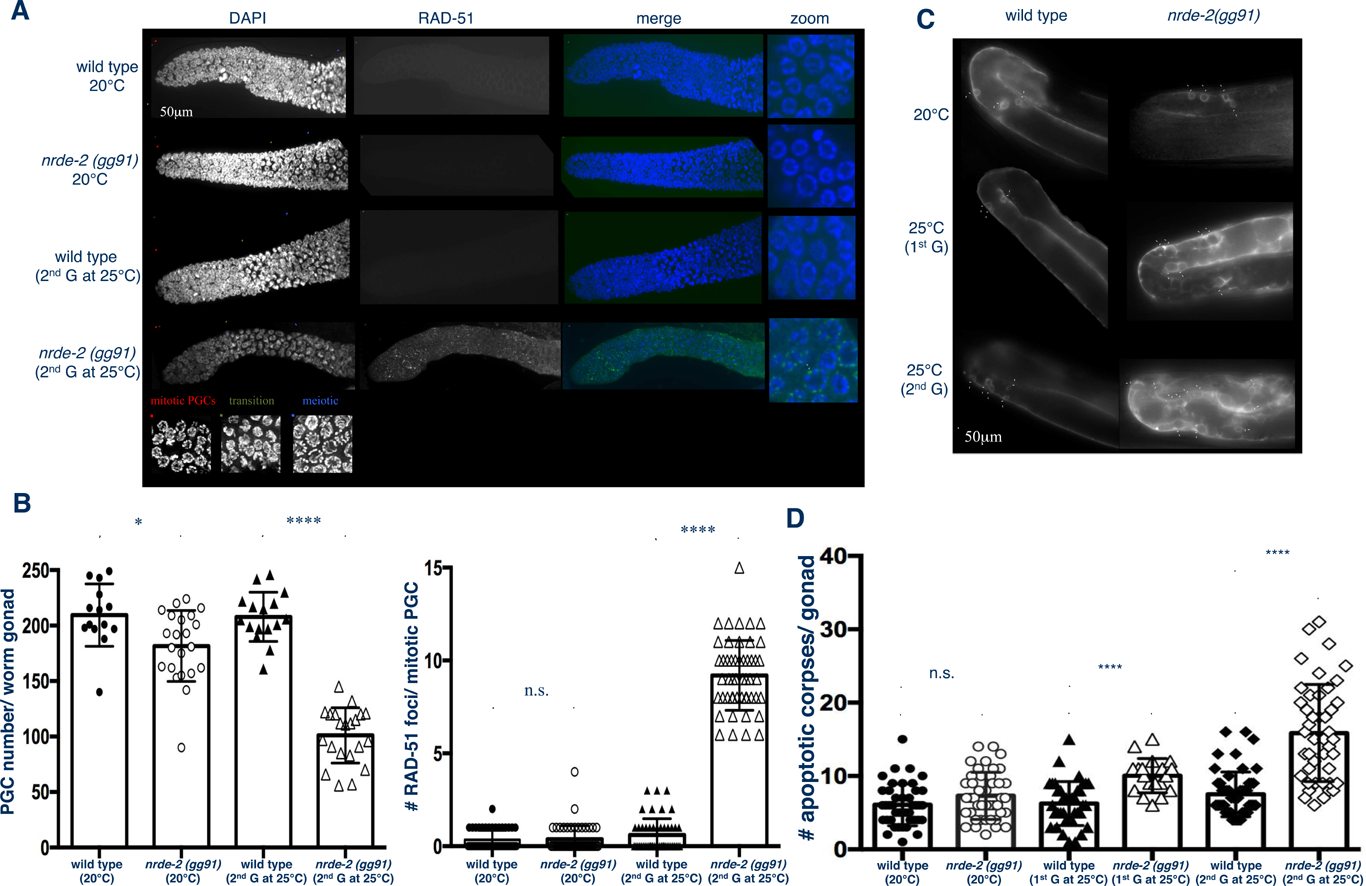
A: Loss of NRDE-2 significantly decreases the number of mitotic germ cells and increases the accumulation of RAD-51 foci in these lls in worms grown for two generations at 25°C. Maximum intensity projections of DAPI stained gonads with RAD-51 foci visualized in the green hannel using an antibody against RAD-51. (Left panels) The red line indicates mitotic proliferating germ cell zone, the green line indicates transition one where DAPI labeled chromosomes exhibit a “crescent-like” morphology, and the blue line indicates meiotic zone. (Right panels) Zoomed in image of one focal plane of mitotic PGCs. The white arrow indicates a single RAD-51puncta. **Figure 1B: (Left Panel) Loss of NRDE-2 at 20°C decreases the number of PGCs and this is exacerbated when the worms are grown for two generations at 25°C.** Mean number + SEM of PCGs in gonads of wild type propagated at 20°C (209.6 ± 7.518, n=14) and *nrde-2(gg91)* mutants propagated at 20°C (181.6 ± 6.830, n=22) are different (*p=0.0115). PGCs of G2 wild type worms propagated at 25°C (207.9 ± 5.542, n=16) and G2 *nrde-2(gg91)* mutants propagated at 25°C (101.0 ± 5.418, n=21) is different (****p<0.0001). **(Right panel) Loss of NRDE2 at 25°C increases the number of RAD-51 foci per PGC nucleus.** The focal plane with the greatest number of RAD-51 foci was chosen for each germ cell nucleus. The number of RAD-51 foci per nuclear focal plane cell were averaged across 50 mitotic PGCs (10 PGCs closest to the distal tip cell in 5 gonads per condition). Mean number + SEM of RAD-51 foci in gonads of wild type propagated at 20°C (0.3400 ± 0.07346, n=50) and *nrde-2(gg91)* mutants propagated at 20°C (0.3800 ± 0.1026, n=50) are not different. RAD-51 foci of G2 wild type worms propagated at 25°C (0.6000 ± 0.1245, n=50) have more RAD-51 foci than G2 *nrde-2(gg91)* mutants propagated at 25°C (9.200 ± 0.2665, n=50, ****p<0.0001). All data represents 3 experimental days and the p-values were determined using unpaired Student’s t test. **Figure 1C: Loss of NRDE-2 increases the number of CED-1::GFP-labeled cell corpses in worms grown for 1 or 2 generations at 25°C.** CED-1::GFP xpressed in *nrde-2(gg91)* and wild type *C. elegans* gonads (white arrows indicate apoptotic cell corpses). **Figure 1D: Loss of NRDE-2 increases the number of apoptotic corpses in the gonads of worms grown for 1 or 2 generations at 25°C.** Quantification of CED-1::GFP labeled germ cells. Wild ype worms propagated at 20°C (6.071 ± 0.4409, n=42) have the same number of corpses as *nrde-2(gg91) mutants propogated* at 20°C (7.302 ± 0.4897, n=43). Wild type worms (6.263 ± 0.4869, n=38) have fewer cell corpses than *nrde-2(gg91)* mutants when grown for 1 generation at 25°C (10.05 ± 0.5379, n=19, ****p<0.0001). Wild type worms (7.510 ± 0.4278, n=51) have fewer cell corpses than *nrde-2 (gg91*) mutants when grown for 2 generations at 25°C (15.86 ± 0.9987, n=44, ****p<0.0001). Cell corpses identified as described in Figure 1D. Data represents 3 experimental days and he p-values were determined using unpaired Student’s t test.

Since the Mrt phenotype in *nrde-2 (gg91)* mutants depends on cultivation at 25°C, we asked whether the cells accumulated DNA damage at this temperature, which would ultimately cause sterility. We used immunofluorescence to stain for RAD-51, a marker of DNA damage repair, and found that *nrde-2 (gg91)* G2 worms grown at 25°C had significantly increased RAD-51 foci in the mitotic zone as compared to wild type G2 worms grown at the same temperature (Fig 1A and B, right, p<0.0001). We then asked whether the aberrant accumulation of DNA damage in the mitotic PGCs resulted in defects as the cells entered meiosis in the proximal gonad and progressed towards oocyte formation. We expressed the apoptotic cell corpse marker, CED-1::GFP, which labels apoptotic cell corpses in the proximal loop of the gonad, in *nrde-2 (gg91)* mutants and wild type worms. We found that when the worms were propagated at 20°C, *nrde-2 (gg91)* mutants showed a similar number of CED-1::GFP labeled germ cell corpses as wild type worms. However, *nrde-2 (gg91)* G1 and G2 worms showed increased numbers of CED-1::GFP labeled germ cell corpses as compared to wild type G1 and G2 worms, respectively (Figure 1C and D, p<0.0001 for both).

We conclude that when worms are cultivated at a normal, non-stressful temperature, loss of NRDE-2 has minor endurable impacts on fertility and integrity of the *C. elegans* germ line. However, when NRDE-2 loss is coupled with an external stress, such as a high cultivation temperature, the DNA damage and subsequent cell death in the gonad compounds over the generations, ultimately resulting in sterility. Therefore, we propose the NRDE-2 may have a role in preventing DNA damage accumulation in the mitotic PGCs in order to maintain integrity of the *C. elegans* germ line.

### Loss of NRDE2 coupled with Aurora B Kinase overexpression in MCF10A cells results in DNA damage accumulation

To gain a biochemical and cell biological understanding of the role of NRDE-2, we decided to utilize the actively dividing MCF10A cell line. MCF10A cell are a normal human epithelial cell line derived from reduction mammoplasty. [4]

We used a lentiviral vector to stably express an eGFP-tagged version of human NRDE2 in the MCF10A cells. The eGFP::NRDE2 localized to puncta within the nucleus (Fig 2Ai). These puncta were reminiscent of staining of heterochromatin and heterochromatin associated factors. [49] Once the cell entered prophase, as characterized by chromatin condensation, the eGFP signal became diffuse and remained so throughout mitosis (Fig 2Aii-v) until cytokinesis, when eGFP::NRDE2 seemed to re-localize to puncta in the nuclei of the daughter cells (Fig 2Avi). The dynamics of this pattern of staining is similar to that of human RNA pol II transcription machinery proteins TBP (TATA box-binding protein) and TAF12 (TATA box-binding protein associated factor 12) in HeLa cells. [29] These data are therefore consistent with a conserved role for NRDE2 as a transcriptional regulator in human cells.

**Figure 2.**
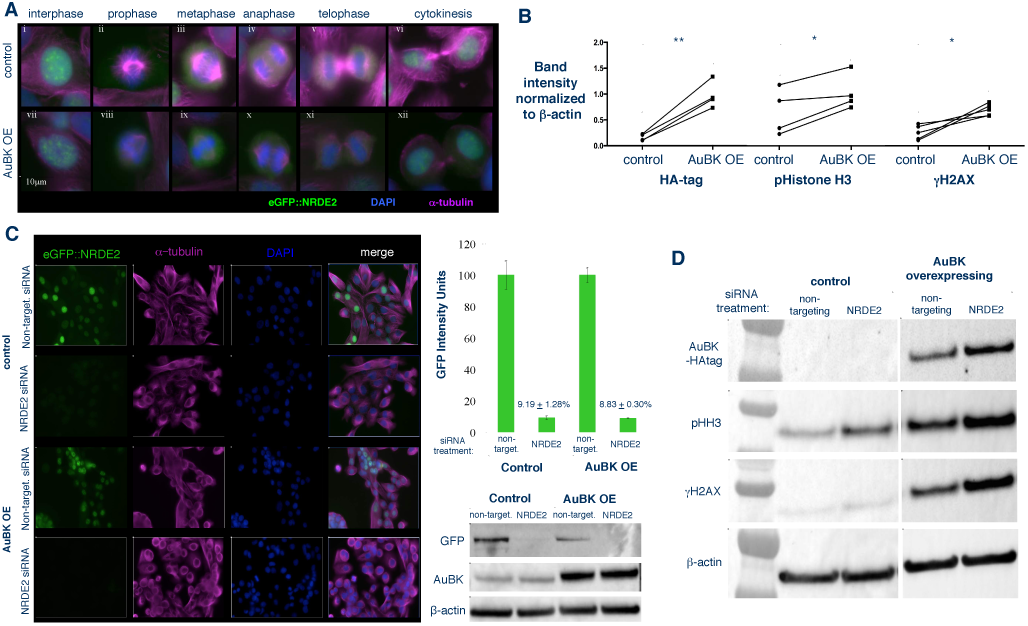
A: In MCF10A human breast cells with endogenous or overexpressed AuBK levels, eGFP-NRDE2 localizes to puncta within the nucleus when chromatin is de-condensed during interphase and cytokinesis. In each circumstance, eGFP-NRDE2 is diffuse throughout all other stages of mitosis. Epifluorescent images of cells expressing eGFP-tagged NRDE2 as visualized in the green channel, chromatin as visualized by DAPI staining in the blue and the cytoskeleton as visualized by an antibody against α-tubulin in the Cy5 (far red) channel. (Top row) MCF10A cells expressing control vector. (Bottom row) MCF10A cells over-expressing AuBK. **Figure 2B: AuBK overexpression in MCF10A cells as a means to stress cells by increasing basal levels of DNA damage.** Western blot band intensities (see figure 2D for representative blot) were measured and normalized to β-actin band intensity. HA-tag band intensity in HA-tagged AUBK overexpressing cells is higher than in control cells (n=4; p=0.0046). Phosphorylated Histone H3 band intensity in AUBK overexpressing cells is higher than in control cells (n=4; p=0.0337). Phosphorylated H2AX band intensity in AUBK overexpressing cells is higher than in control cells (n=5; p=0.0146). All p-values determined by paired Student’s t test. **Figure 2C: Average GFP intensity in MCF10A cells expressing eGFP-tagged NRDE2 decreases specifically in cells treated with NRDE2 targeted siRNA.** (Left) Epifluorescent images of MCF10A cells expressing eGFP-tagged NRDE2 as visualized in the green channel, the cytoskeleton as visualized by an antibody against α-tubulin in the far red channel and DNA as visualized by DAPI staining in the blue channel. Control or AuBK overexpressing MCF10A cells were treated with non-targeting siRNA or NRDE2 siRNA. (Top right) Quantification of epifluorescent GFP intensity of control and AuBK overexpressing cells treated with NRDE2 siRNA and normalized to the respective non-targeting siRNA treated cells. (Bottom right) Representative western blot using anti-GFP to detect eGFP-tagged NRDE2 inMCF10A cells treated with non-targeting or NRDE2 targeted siRNA. All cells treated with siRNA for 72 hours before imaging. Data represents 4 experimental days. **Figure 2D: MCF10A cells that overexpress HA-tagged AuBK show an increase in phosphorylated Histone H3 and an increase in the DNA damage marker γH2AX.** Representative blot: Anti-HA-tag used to detect AuBK in cells transformed with AuBK-HAtag in MCF10A cells. AuBK overexpressing cells have increased levels of phosphorylated Histone H3 and increased levels of γH2AX expression as compared to control cells. Western blot band intensities were measured and normalized to β-actin band intensity. See Fig S3 for quantification and statistics.

We wanted to further test the hypothesis that NRDE-2 regulates DNA damage and mitosis under stressful conditions. We therefore created a mitotically stressed MCF10A cell line by stably overexpressing an HA-tagged version of the oncogene Aurora B kinase (AuBK OE) in eGFP::NRDE-2 expressing cells. AuBK is a master regulator of mitotic events, including chromatin condensation during prophase, [5] microtubule-kinetochore attachment during metaphase, [6], [7] and abscission during cytokinesis. [8] Loss of this regulation has been linked to genetic instability, which results in aneuploidy, a key hallmark of cancer. Accordingly, AuBK over-expression has been linked to many cancer types and correlates with poor prognosis for patients. [11–12] The localization of eGFP::NRDE2 in AuBK OE MCF10A cell lines throughout the cell cycle was identical to that of eGFP::NRDE2 in control cells (Fig 2A vii and xii).

Consistent with AuBK’s role in mediating chromatin condensation at the G2/M transition, AuBK OE MCF10A cells showed increased levels of phosphorylated Histone H3 (Fig 2B, middle, p=0.0337, n=4), a known target of Aurora B Kinase. We also observed a significant increase in γH2AX (H2AX phosphorylation on S139), a common marker of DNA damage (Fig 2B, right, p=0.015, n=5). Therefore, we utilized AuBK overexpression as a tool to study the effects of NRDE2 loss on DNA damaged cells. Next, we assessed the efficacy of using NRDE2 targeted siRNA as a tool to knockdown NRDE2 in MCF10A cells. We observed that cells treated with NRDE2 siRNA showed >90% decrease in eGFP::NRDE2 expression as compared to cells treated with a non-targeting siRNA control (Fig 2C). We then asked whether the amount of DNA damage in control or AuBK OE cells was altered upon NRDE2 knockdown. Interestingly, we found when control cells were treated with NRDE2 siRNA, there was no increase in γH2AX as compared to control cells treated with non-targeting siRNA (Fig S3, p=0.11, n=5). However, loss of NRDE2 in AuBK OE cells resulted in increased DNA damage as compared to AuBK OE cells treated with non-targeting siRNA (Fig 2D, p=0.048, n=5). We conclude that loss of NRDE2 has a minimal effect on DNA damage in normally proliferating cells. However, in AuBK OE cells, which show an increased basal level of DNA damage, loss of NRDE2 further increases the accumulation of DNA damage.

### Loss of NRDE2 coupled with Aurora B Kinase overexpression in MCF10A cells results in mitotic delay

We asked whether the increase in DNA damage observed in the AuBK OE cells upon NRDE2 loss would result in an observable change in mitotic cell proliferation. We observed that when AUBK OE cells were treated with NRDE2 siRNA, there was a decrease in overall metabolic cell activity in the culture (Fig S2) and a significant increase in the number of rounded cells (Fig 3A). Cells commonly round up due to a stall in mitosis or when the cell is about to undergo apoptosis, both of which could explain the observed decrease in metabolic cell activity. To differentiate between these two possibilities, we used live cell imaging to record cell growth after NRDE2 knockdown. We found that AuBK OE cells treated with non-targeting siRNA took a longer time to complete mitosis than control cells. Furthermore, when AuBK OE cells were treated with NRDE2 siRNA, there was a significant increase in the average time through mitosis (Fig 3B and C). However, we also noticed that a small proportion of AUBK-overexpressing cells required a significantly longer time to complete mitosis upon NRDE2 siRNA treatment (Fig 3B and C). The rounded morphology of the cells, the compaction of the chromatin and the concentration of the DNA at the cell center, indicated that this stall may occur during the early stages of mitosis.

**Figure 3.**
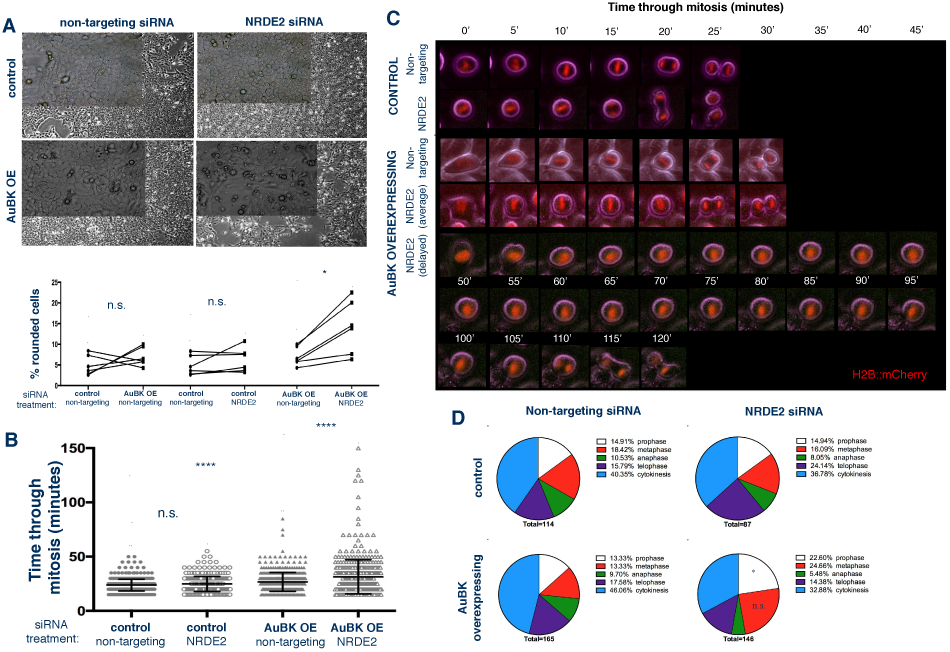
A: Loss of NRDE2 in MCF10A cells overexpressing AuBK results in an increase in the percentage of cells with a rounded morphology. (Top) Brightfield images of cells 72 hours after siRNA treatment taken with 4x objective and 20x objective (inset). (Bottom) Quantitation of cell morphology. (Left paired points) Control or AuBK OE MCF10A cells show no difference in the percentage of round cells in cultures treated with nonargeting siRNA. (Middle paired points) MCF10A control cells show no difference in the percentage of round cells upon NRDE2 siRNA treatment as ompared to non-targeting siRNA treatment. (Right paired points) MCF10A AuBK OE cells show a significant increase (*p=0.0110) in the percentage of round cells upon NRDE2 siRNA treatment as compared to non-targeting siRNA treatment. Data represents 6 experimental days. All p-values determined by paired Student’s t test. **Figure 3B: Loss of NRDE2 in MCF10A cells overexpressing AuBK delays time through mitosis.** Time required for individual MCF10A cells to through mitosis. Control cells treated with NRDE2 siRNA (24.87 ± 0.3902, n=318) do not take significantly onger than control cells treated with non-targeting siRNA (23.79 ± 0.2442, n=492) to progrss through mitosis (p=not significant, Mann-Whitney test). AuBK over-expressing cells treated with NRDE2 siRNA (31.40 ± 0.7220, n=472) take significantly longer than cell trated with non-targeting siRNA 26.72 ± 0.4120, n=451) to progress through mitosis (****p<0.0001, Mann-Whitney test) **Figure 3C: Loss of NRDE delays time through mitosis in a subset of MCF10A cells overexpressing AuBK** of live MCF10A cells expressing H2B::mCherry visualized in the red channel to visualize hromatin compaction and chromosome separation. **Figure 3D: Loss of NRDE2 in AuBK overexpressing MCF10A cells significantly increases the proportion of cells in prophase.** Quantification of the percentage of cells in each phase of mitosis shows that AuBK overexpressing MCF10A cells have significantly more cells in prophase after treatment with NRDE2 siRNA as compared to treatment with non-targeting siRNA (p= 0.0258, determined by student’s t test). Data represents 4 experimental days.

We propose that the rounded cells observed upon NRDE2 knockdown in AUBK-overexpressing cells is due to a delay in mitosis. To further characterize this delay, we calculated the mitotic index of these cells. Consistent with the time lapse data, we found that AuBK OE cells had a higher mitotic index than control cells. However, we found no significant difference between the mitotic index of non-targeting siRNA treated cells and the NRDE2 siRNA treated cells in either the control or AuBK OE (Fig S4). We surmised that this could be due to the fact that the percentage of cells with a significant delay in mitosis is so low that these cells were not sufficient to affect the mitotic index.

We next asked whether we could identify the stage at which mitosis was delayed in AUBK-OE cells. We quantified the number of cells in each stage of mitosis in a fixed population of cells and found more cells in prophase upon NRDE2 knock down (Fig 3E p=0.025, n=4). This is also consistent with the time lapse analysis of AuBK OE cells delayed in early mitosis (Fig 3C) leading us to conclude that the delay seen in NRDE2 knockdown AUBK-overexpressing cells may be occurring at prophase.

Mammalian NRDE2 interacts with β-actin in healthy, normally proliferating cells, and this interaction is weakened upon DNA damage Our finding that NRDE2 specifically mitigates DNA damage accumulation in sensitized or stressed cell populations led us to ask whether NRDE2 interacts with different partners under these conditions. We decided to probe this question using coimmunoprecipitation experiments, using an anti-GFP antibody to detect eGFP::NRDE2 in MCF10A lysates (Fig 4A and Fig S5).

**Figure 4.**
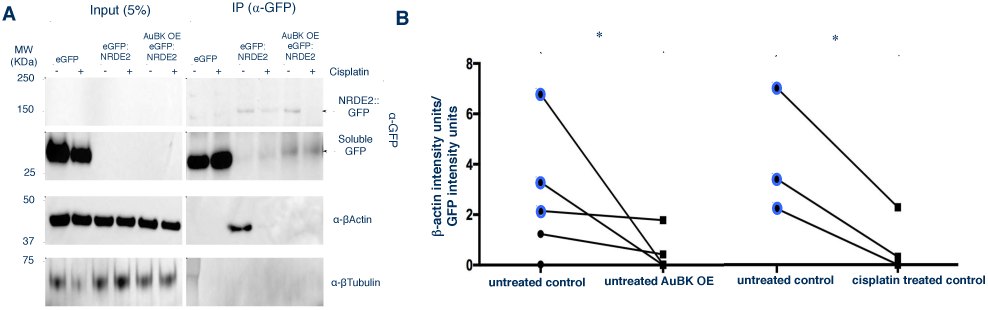
A: eGFP::NRDE2 interacts with β-actin in control cells. This interaction is weakened in AuBK OE cells and when cells are treated with cisplatin. Representative western blots of MCF10A cell lysates. MCF10A cells were treated with 60uM cisplatin for 16 hours. Co-IP was performed using goat Anti-GFP antibody to pull down eGFP::NRDE2 from MCF10A cell lysates. Western blots were developed using rabbit anti-GFP polyclonal antibody to detect eGFP::NRDE2 and soluble GFP, anti-β-actin HRP antibody and mouse anti-β-tubulin antibody (negative control). **Figure 4B: The amount of β-actin per unit of IP’d GFP::NRDE2 decreases when DNA damage is induced in cells by AuBK OE or by cisplatin drug treatment.** (Left) The amount of β-actin per unit of co-IP’d eGFP::NRDE2 decreases when MCF10A cells overexpress AuBK as compared to MCF10A control cells (*Permutation test exact p-value =0.036). (Right) The amount of β-actin per unit of co-IP’d eGFP::NRDE2 decreases when MCF10A control cells are treated with 60uM cisplatin as compared to untreated control MCF10A cells. (*p-value =0.045 as assessed by paired student’s t test). Note: Blue data points represent quantification of the same samples.

In *C. elegans*, NRDE-2 interacts with other nuclear RNAi factors NRDE-1 and NRDE-4, as well as the worm specific argonaute NRDE-3. [11] In contrast, NRDE2 does not interact with other RNAi factors in S. pombe yeast cells or in human cells. [22, 43, 44] Consistent with these findings, we saw no interaction between eGFP::NRDE2 and any of the known human Argonautes AGO-1/2/3/ or ‐4 (Fig S6).

Interestingly, we observed a strong interaction between NRDE2 and β-actin in control cells (Fig 4A). Furthermore, this interaction seemed to be weakened in AuBK OE cells (Fig 4A and B, p=0.036, n=5). These results may suggest an interesting function for NRDE2 in DNA damaged cells. Nuclear actin forms filaments upon DNA damage to aid in repair. [25] Since AuBK overexpression increases DNA damage in MCF10A cells, we hypothesized that the interaction between NRDE2 and β-actin may be weakened in these cells due to the nuclear actin forming filaments in response to DNA damage. To test whether DNA damage altered the interaction between NRDE2 and β-actin, we treated the cells with cisplatin, a DNA damaging reagent. In both control and AuBK overexpressing MCF10A cells, the interaction between NRDE2 and β-actin was reduced as compared to untreated cells (Fig 4A and B, p=0.045, n=3). Therefore, we propose that NRDE2 interacts with β-actin only when the DNA is intact. However, when DNA damage in induced by stressors such as AuBK over-expression or the presence of cisplatin, the interaction between NRDE2 and β-actin is disrupted, perhaps as a result of the nuclear actin forming filaments to aid in DNA repair.

## Discussion

This study describes a previously uncharacterized role for NRDE2, a putative nuclear RNA-binding protein, in protecting against DNA damage in stressed cells. Our study is the first to study the effect of NRDE2 loss in normal, unstressed cells as compared to in stressed cells. We have also shown that this protective role for NRDE2 in stressed cells is conserved in *C. elegans* and mammalian mitotic cells.

Previously, NRDE2 was described to protect against the accumulation of DNA damage, as assessed by an increase in γH2AX levels upon NRDE2 loss in HEK293 cells. [22] We believe this is consistent with our results as HEK293 cells are not a normally proliferating cell line. The HEK293 line was originally obtained by transforming human embryonic kidney cells with human adenovirus type 5, resulting in a 4.5kb insertion of a viral fragment in chromosome 19 which interferes with cell death and cell cycle control pathways. [23] HEK293 cells are also hypotriploid, [24] which may further affect mitosis. Consistent with this point, we found that NRDE2 knockdown resulted in an increase of γH2AX levels only in AuBK-overexpressing MCF10A cells, not in the normal, unstressed control cells. Since AuBK overexpressing causes an increase in basal levels of DNA damage, we propose that NRDE2 in mammalian cells results in an accumulation of DNA damage only in cells with increases mitotic stress, similar to the results seen in *C. elegans*.

In *C. elegans*, loss of NRDE-2 alone does not seem to severely compromise the mitotic PGCs or brood size, as evidenced by the worms remaining indefinitely fertile and exhibiting relatively normal development. Upon the addition of a stressful 25°C cultivation temperature, however, there is a drastic reduction in the number of PGCs and brood size. Under these stressful conditions, DNA damage accumulates in the germ cells, presumably resulting in the previously observed Mrt phenotype. This trans-generational Mrt phenotype is similar to fertility defects observed *rsd-2* and *rsd-6* RNAi spreading defective mutants. RSD-2 and RSD-6, along with NRDE-2 were proposed to induce genome wide epigenetic silencing to maintain germ cell immortality and fertility. RSD-2 and RSD-6 are also thought to act in a different pathway than the 22G-RNA associating argonautes CSR-1 and ALG3/4, which display Mrt phenotypes due to defects in spermatogenesis. [16] The effect of NRDE-2 loss on spermatogenesis remains unclear.

The defects seen in *nrde-2 (gg91)* mutants may also be due to improper chromatin condensation and chromosome segregation. It has been previously reported that CSR-1 is required for the localization of CDE-1, a nucleotidyltransfease protein that is responsible for the uridylation of siRNAs bound by CSR-1, to mitotic chromosomes in embryos. [19] Loss of CDE-1 results in an accumulation of siRNAs and improper gene silencing, as well as mitotic and meiotic chromosome segregation defects. NRDE-2, by contrast, associates with other NRDE pathway proteins to recruit methlytransferases MES-2 and SET-25, which direct the deposition of methylation marks to silence loci targeted by siRNAs. [20–21] Therefore, NRDE-2 loss may also result in improper gene silencing in the *C. elegans* germ line, resulting in mitotic defects, which are exacerbated by the stress of high temperature.

Interestingly, we observed a strong association between NRDE2 and β-actin in MCF10A cells that was diminished upon DNA damage causing agents. Since NRDE2 localizes to puncta in the interphase nucleus, we hypothesize that NRDE2 likely interacts with nuclear actin. One potential model for this interaction is that NRDE2 binds to monomeric nuclear actin in the unstressed cell. Monomeric nuclear actin associates with chromatin remodeling complexes, and might be involved in chromatin binding. [45] However, upon DNA damage, nuclear actin assembles into filaments and promotes DNA damage repair. [25] NRDE2 may be displaced or unable to bind to filamentous actin, and as a result, show a weakened interaction on β-actin upon DNA damage. It is still unclear whether NRDE-2 functions in preventing the formation of DNA damage or as a chromatin associated sensor of DNA damage in the β-actin dependent DNA repair pathway. Further study is needed to characterize the function of NRDE2, and possibly other heterochromatin factors, in preventing DNA damage accumulation and maintaining genomic stability. The interaction between NRDE2 and β-actin is especially exciting as it may indicate a new mechanistic role for NRDE2 in the nucleus beyond gene silencing and epigenetic inheritance.

## Materials and Methods

### *C. elegans* strains

All strains were cultured on Nematode Growth Medium (NGM) plates seeded with E. coli OP50. Bristol N2 was the wild type strain used. Loss of NRDE-2 was studied using the mutant allele *nrde-2 (gg91).* Worms were crossed into a strain expressing ced-1p::GFP+ lin-15 (+) to study apoptotic cells.

### Gonad Staining

Gonads were dissected from young adults (24 hours post-L4 stage) in 1xEGG buffer (25mM HEPES, pH7.3; 118mM NaCl, 48mM KCl, 2mM CaCl2, 2mM MgCl2) and fixed in 2% (final concentration) formaldehyde. The following primary antibodies were used: rabbit anti-RAD-51 and chicken anti-HTP-3. Fixed gonads were imaged with exposure times of 100ms with DAPI, GFP, TRITC, and CY5 filter cubes and a mercury arc lamp on a Zeiss AxioPlan2 epifluorescence microscope (operated by MicroManager 1.4.13) with a 60x 1.3 DIC oil objective and a QIClick camera (QImaging). Images were recorded with a Z-optical spacing of 2 μm and analyzed using the Image J software.

### Progeny Assay

Sixteen L4-staged worms were individually placed on a 5cm NGM plate with op50. Every 24 hours, the worm was moved to a fresh plate for 10 consecutive days. 48 hours after the original worm was removed from the plate, the number of living progeny was counted.

### Cell Culture

MCF10A cells lines expressed a vector conferring puromycin/ blastocydin antibiotic resistance that was empty (control) or that drove the overexpression of Aurora B Kinase (AuBK OE). MCF10A cells were maintained in Dulbecco’s modified Eagle’s medium/F-12, phenol red free medium supplemented with 5% heat inactivated horse serum, 20 ng/ml recombinant EGF (Invitrogen), 0.5 μg/ml hydrocortisone (Sigma), 100 ng/ml cholera toxin (Sigma), and 10 μg/ml insulin (Sigma). U20S cells were maintained using Dulbecco’s modified Eagle’s medium with phenol red supplemented with 10% FBS.

### siRNA knockdown

We used the following Thermo Scientific siRNAs: siGENOME Non-Targeting siRNA

Pool#2 (D-001206-14-20), siGENOME Human NRDE2 (55051) siRNA‐ SMARTpool (M-013794-00-0005). siRNAs were added to the cell culture at a final concentration of 5nM in OPTIMEM Reduced Serum Media with Lipofectamine 2000 Reagent (1ug/ml final concentration) (Invitrogen) for 24 hours. Cells were then washed 1x with PBS and fresh MCF10A media was added. 48 hours after changing media, the cells were harvested for analysis.

### Western Blot

Cultured cells were homogenized in Laemmli Buffer (60 mM Tris–HCl pH 7.6, 2% SDS, 1mM DTT) containing COMPLETE protease inhibitor cocktail (Roche #118361700001) and phosphatase inhibitors (Santa Cruz Biotechnology)). Protein concentrations were determined by performing DC Protein Assay (Bio-Rad) using BSA as standard. Protein extracts were resolved using Bolt 4%-12% Bis-Tris PAGE gels (Invitrogen) with 1x MES SDS Running Buffer (Life Technologies). Transfer to nitrocellulose membranes (Life Technologies) was performed on an iBlot apparatus (Invitrogen). Membranes were probed with primary antibodies overnight on a 4°C shaker and then incubated with horseradish peroxidase (HRP)-conjugated secondary antibodies, and signals were visualized with ECL (Bio-Rad) or Visualizer Western Blot Detection Kit, rabbit (Millipore). The following primary antibodies were used: GFP (Rockland, 600-101-215), phospho-Histone H3 (Ser10) (Cell Signaling Technologies, #9701), phospho-Histone H2A.X (Ser 139) (Cell Signaling Technologies, #9718), HA-tag (Cell Signaling Technologies #2367), PARP (Cell Signaling Technologies, #9542), β-actin HRP (Santa Cruz Biotechnology, sc-47778)

### Cell Immunofluorescence

Cells were grown on a sterilized glass coverslip (Fisherbrand). After siRNA treatment, the cells were fixed with ice cold 100% methanol for 3 minutes at ‐20°C and rehydrated with TBST (1xTBS, 0.1% Tween 20). Antibodies were made up in blocking solution (2gBSA powder, 0.1g NaN3, 100mL TBST). The following primary antibodies were used: human anti-centromere (CREST, 1:50, Antibodies Incorporated 15-234) mouse anti-a-tubulin (DM1a, 1:1000, Sigma T6199), rabbit anti-g-tubulin (1:1000, Sigma T3559). Cells were mounted with Vectashield Mounting Medium with DAPI (Vector Laboratories H-1200).

### Time lapse Imaging

MCF10A cells expressing H2B::mCherry were imaged at 37°C with a 100x NA 1.49 objective lens (CFI APO TIRF; Nikon) on an inverted microscope system (TE2000 Perfect Focus System; Nikon) equipped with a Borealis modified spinning disk confocal unit (CSU10; Yokogawa) with 200-mW, 405 nm, 488 nm, 561 nm and 643 nm solid-state lasers (LMM5; Spectral Applied Research), electronic shutters, a Clara cooled scientific-grade interline CCD camera (Andor), and controlled by NIS-Elements software (Nikon).

### Co-Immunoprecipitation

Cells were collected and lysed in cold lysis buffer (150mM NaCl/ 1.0% IGEPAL/ 50mM Tris-Cl (pH 7.4)) containing Complete protease inhibitor cocktail (Roche, 118361700001) by nutating at 4°C for 30 minutes. The lysate was spun at 14K G, and supernatant was transferred to a new tube, and cleared using 50ul of Dynabeads Protein A (Invitrogen, 10001D). Pre-cleared lysate was removed from the beads using magnetic separation (input sample was collected at this stage), added to 5 ul of anti-GFP antibody (Rockland, 600-101-215), and nutated at 4°C for 2 hours. Antibody-protein complexes were isolated by the addition of Dynabeads Protein A (Invitrogen, 10001D), overnight nutation and magnetic separation. The beads were washed with the lysis buffer containing protease inhibitors five times. IP western blot bands were normalized to 5% sample input.

## Acknowledgements

Thanks to Dr. Scott Kennedy for providing the nrde-2 (gg91) strain and to Dr. Needhi Bhalla (UC Santa Cruz) for providing the CED-1::GFP strain. We would like to thank Dr. Gina Caldas (Dernburg lab at UC Berkeley) for reagents and advice about *C. elegans* gonad immunofluorescence staining. We extend a special thanks to Christina Hueschen (Dumont lab at UCSF) for creating a nuclear GFP quantification MatLab script. We would also like to thank Dr. Sophie Dumont and Dr. Torsten Wittmann (UCSF) for use of their microscopes and advice regarding imaging and the cell cycle.

---

**Figure S1.**
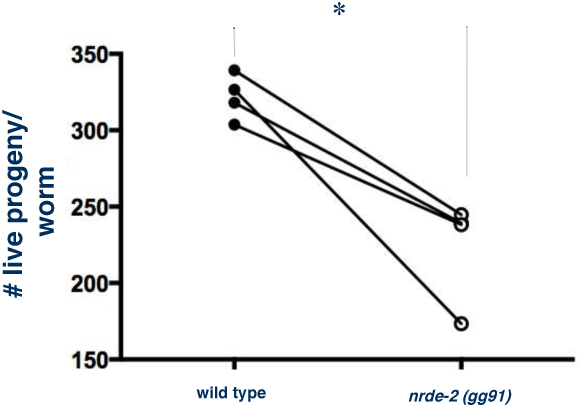
Loss of NRDE-2 significantly decreases brood size as compared to wild type worms when propagated at 20°C. Each data point represents one biological replicate of 16 worms per experiment (n=4; *p=0.0148, paired Student’s t test).

**Figure S2.**
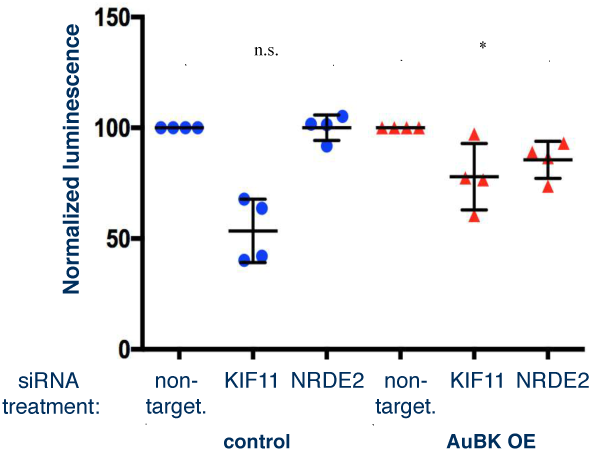
NRDE2 knockdown decreases cell viability in AuBK OE cells but not in control cells. The Cell Titer Glo assay measures the number of metabolically active cells by quantifying the amount of ATP present in the cell culture. The luminescent signal was normalized to non-targeting siRNA treated cells in both control cells and AuBK OE cells. KIF11 siRNA was used as a positive control. Control cells treated with NRDE2 siRNA (100.0 ± 2.890, n=4) had no change in luminescence as compared to control cells treated with non-targeting siRNA. AuBK OE cells treated with NRDE2 siRNA (85.55 + 4.167, n=4) had a lower amount of luminescence as compared to AuBK OE cells treated with non-targeting siRNA (*p=0.0133). P-values determined using unpaired Student’s t test.

**Figure S3.**
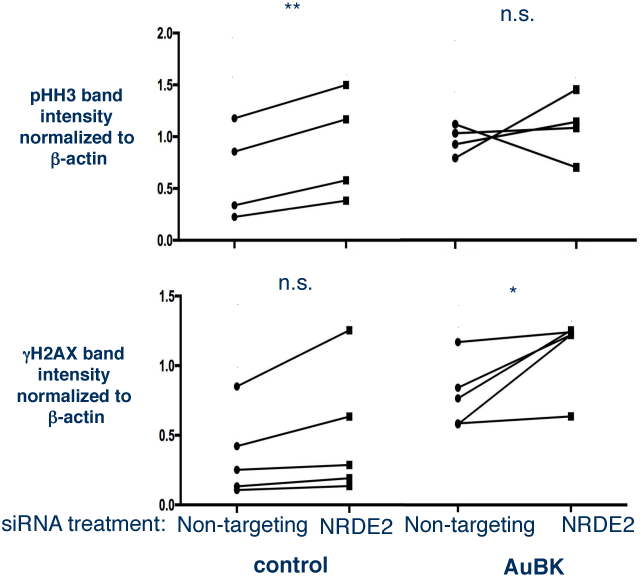
(Top) Loss of NRDE2 in control cells increases pHH3 levels as compared to control cells treated with non-targeting siRNA (n=4, p=0.0068). In contrast, loss of NRDE2 in AuBK OE cells does not change pHH3 levels (p=???). (Bottom) Loss of NRDE2 in control cells does not alter levels of γH2AX (p=????), however in AuBK OE cells, loss of NRDE2 increases γH2AX levels. (n=5, *p=0.0483) All p-values determined using paired student’s t test.

**Figure S4.**
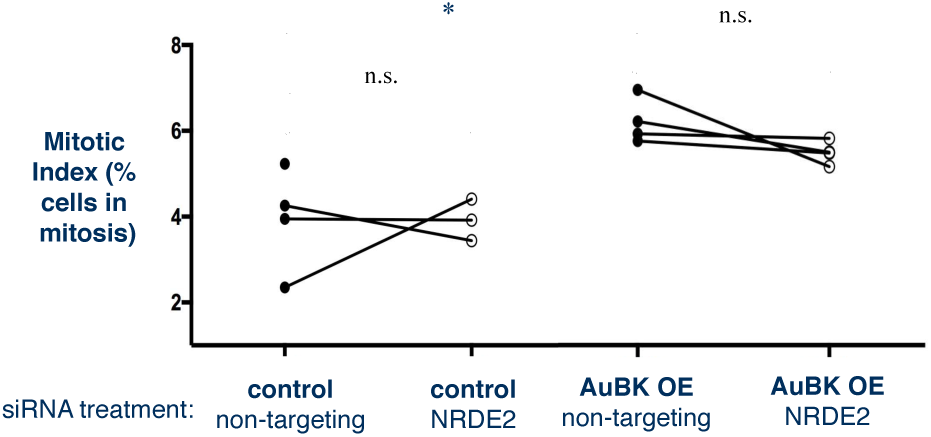
Loss of NRDE2 in MCF10A cells does not significantly change the mitotic index. Mitotic Index calculated by dividing total number of cells in mitosis by total number of cells in the population. AuBK overexpressing cells have a significantly higher basal mitotic index than control cells (*p=0.0315, student’s paired t test).

**Figure S5.**
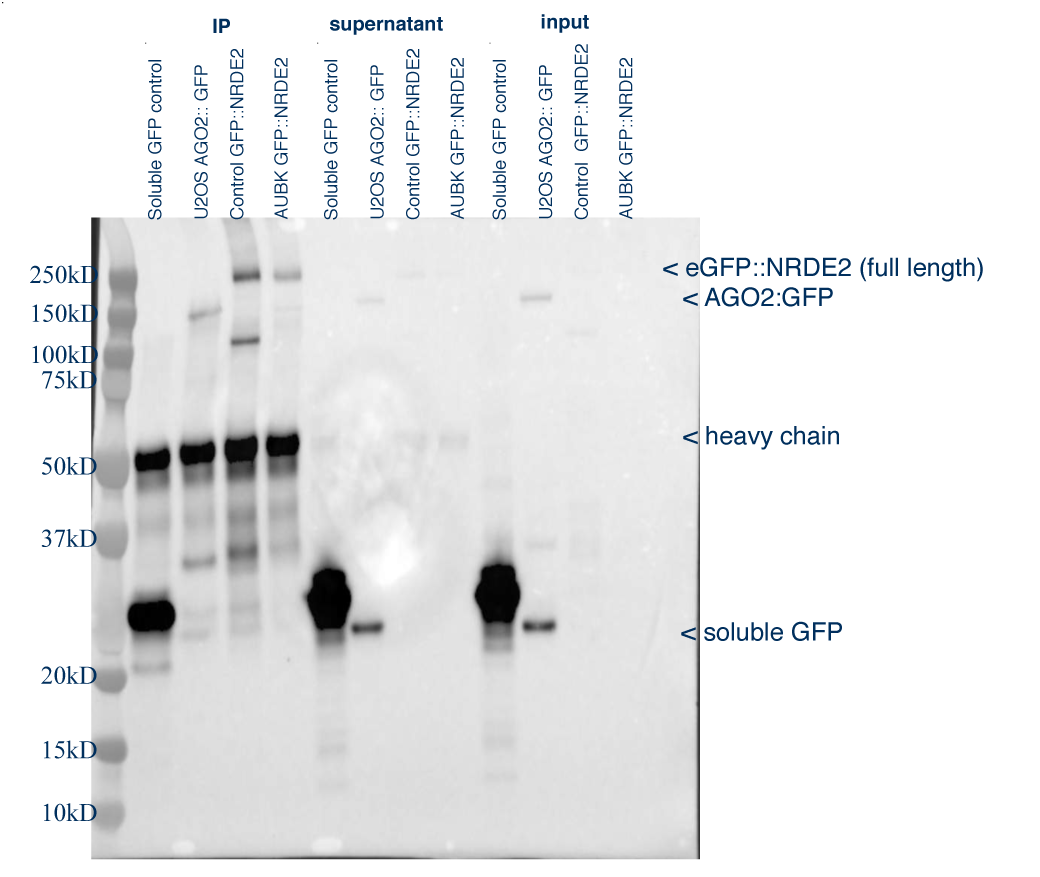
Representative western blot of MCF10A cell lysates. Co-IP was performed using goat Anti-GFP antibody to pull down eGFP::NRDE2 from MCF10A cell lysates. Western blots were developed using rabbit anti-GFP polyclonal antibody to detect eGFP::NRDE2 and soluble GFP.

**Figure S6.**
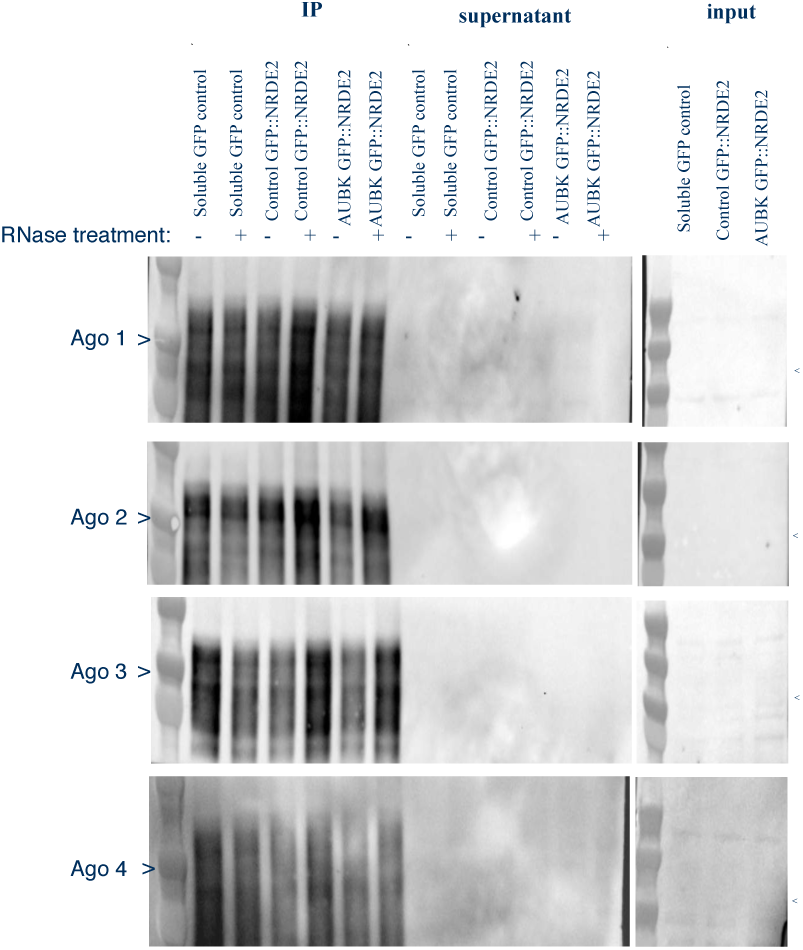
NRDE-2 does not interact with the mammlian argonautes Ago 1,2,3 or 4. Goat anti-GFP used to pull down eGFP::NRDE2 in MCF10A lysates in co-IP assay. Western blots against human Ago1-4 show no interaction with eGFP::NRDE2. Arrows point to 97kD, the expected size of each Ago protein.

**Figure S7.**
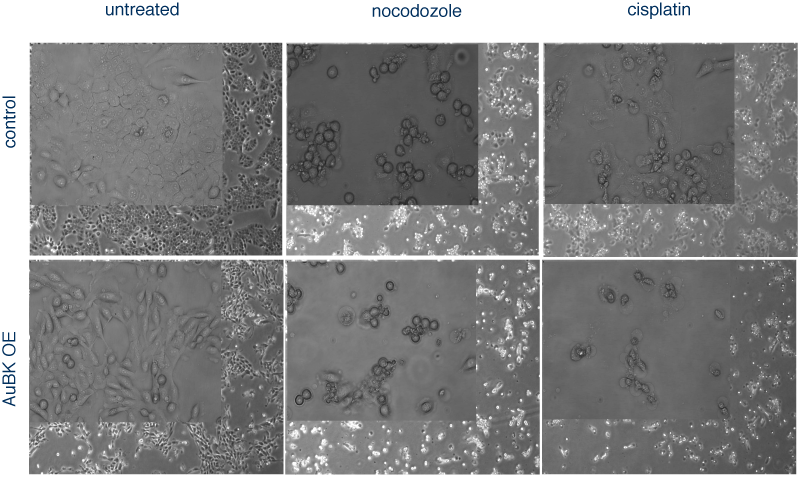
Control and AuBK OE MCF10A cells treated with nocodozole show rounded morphology consistant with stall in G2/M. Cells treated with cisplatin show rounded, fragmented morphology, consistant with apoptosis. Representative brightfield images of cells taken 16 hours after 0.1ug/ml nocodozole or 60uM cisplatin drug treatment with 4x objective and 20x objective (inset).

## Abbreviations

PGC: Proliferative Germ Cell
NRDE-2: Nuclear RNAi Defective-2
AuBK: Aurora B Kinase
Mrt: Mortal germ line

